# Brain and gonadal genes are differentially expressed in *Bicyclus anynana* butterfly larvae that learned a heritable novel food odor preference

**DOI:** 10.1101/2024.08.17.608425

**Authors:** V. Gowri, Shen Tian, Antónia Monteiro

## Abstract

*Bicyclus anynana* butterfly larvae learn to prefer novel odors added to their plant food and transmit those learned preferences to the next generation. However, the molecular mechanisms regulating the inheritance of this acquired preference remain unexplored. Here we examined how larval diet affected gene expression patterns in the larval brain as well as the gonads of adults to explore a potential genetic basis of this inheritance. We fed *B. anynana* larvae leaves coated with a novel banana odor (isoamyl acetate), or with a control solution, and performed five choice assays on individual larvae during their development to identify individuals that showed a majority preference for the treatment odor they were fed with. We then dissected male and female larval brains, adult spermatophores, or adult oocytes from those individuals, and profiled mRNA in all tissues and micro-RNA (miRNA) expression in oocytes only. Our results show that there are 83 differentially expressed genes (DEGs) across all tissue types in odor and control groups, of which 23 play roles in metabolism, transcription, response to various chemical stimuli, and olfactory pathways. MiRNAs did not differ in expression across diet treatments, but we discovered 57 novel miRNAs in oocytes. The DEGs in gonads are potential epigenetic factors that can regulate the inheritance of a learned odor preference. Still, caution is required as there was no overlap between DEGs across male and female brains, as well as male and female gonads.

## Introduction

Recent studies have started to examine the molecular mechanisms underlying the development and inheritance of environmentally acquired characteristics in organisms [1,2]. RNA molecules or epigenetic marks found on the DNA or in histone tails, can regulate gene expression and can also play a crucial role in the transmission of heritable changes that do not involve changes in DNA sequences [3–5]. For instance, odor-conditioned male mice showed differential expression of olfactory receptor genes in their sperm [6], and stress-associated genes were differentially expressed in the brain of their offspring, relative to controls [7]. Small non-coding RNAs, such as micro-RNAs (miRNAs), were involved in the olfactory learning and memory formation via gene regulation at both transcriptional and post-translational levels in mice, fruit flies, rats and honeybees [8–13]. For example, differentially expressed miRNAs that target cell growth regulation and cellular responses were found in the sperm of odor-conditioned male mice [14]. An aversive behavior towards an odor in offspring of nematode worms was inherited via the sperm of conditioned parents, involving the interplay of histone methylation, small RNAs and neuropeptides [15], showing that these miRNAs are epigenetically transmitted. Although these studies focus on changes upon odor exposure paired with either rewards or stressful conditions, not much is known regarding the epigenetic mechanisms or factors that play a role in odor learning via simple odor feeding, and odor preference inheritance.

Here, we tested for differences in mRNA and miRNA expression profiles in brains and in the germline between control-fed and odor-fed larvae and adult butterflies. We used *Bicyclus anynana*, a subtropical model nymphalid butterfly, because this species learns to prefer novel food odors during larval growth [16], or pheromone blends during adult courtship [17], and passes down these learned odor preferences to the next generation [16,18]. In this experiment, we pursued findings from previous studies that show that a single generation of feeding on a novel food odor can induce a preference for that odor in older larvae. Furthermore, when these larvae become adults, they transmit the learned odor preference to their naïve offspring [16,19]. Factors that induce this learned odor preference are found in the hemolymph because odor-fed larvae are capable of inducing an odor preference in the offspring of naïve individuals via mere hemolymph transfusions [20].

In this study, we fed larvae throughout their larval stage with their normal diet of corn leaves, but coated the leaves with a novel chemical odor, or control odor, as in previous experiments [16,19,20]. We selected those larvae that showed a majority preference for their treatment odor. We then performed RNA sequencing of the larval brains, adult oocytes and spermatophores, and small RNA sequencing of the oocytes only. We hypothesized that we might find differences in gene or miRNA expression profiles between both control- and odor-fed groups. Genes differentially expressed in brains might be the same or different from those in the gonads. These differentially expressed RNAs could be epigenetic factors driving the inheritance of the learned odor preference in *B. anynana*.

## Materials and methods

### Husbandry

*B. anynana* were reared in a climate-controlled room at 27°C, 60% humidity, and 12:12-hour light: dark photoperiod. The larvae were fed with corn leaves (*Zea mays*) and the adults were fed on mashed banana. Corn leaves were kept in adult cages for at least three hours to collect eggs. Once the naïve 1^st^ instar larvae hatched they were tested for their innate odor preference (day 0). Larvae that chose control were assigned to a control treatment, those that chose odor were assigned to an odor treatment. Each larva was reared individually in small plastic cups, and they were tested for their odor preferences again on days 5, 10, 15 and 20.

### Preparation of scented leaves

Isoamyl acetate (IAA), also known as isopentyl acetate, from Sigma-Aldrich (W205532, natural, ≥97%, Food Chemicals Codex, Food Grade) was the test odor used. It smells of ripe bananas. A 5% IAA solution was prepared using absolute ethanol (Fisher Chemical, 99.8%, Analytical reagent grade) as IAA is highly insoluble in water. Absolute ethanol was used as the control solution. Corn leaves were coated thoroughly on both sides with the solutions using cotton balls. Ethanol from the coated leaves of both treatments was allowed to evaporate completely before feeding the leaves to larvae. The feed was replaced regularly with freshly coated leaves (in both treatments) to ensure continuous exposure (of odor) to the larvae.

### Odor choice assay

The odor choice made by each larva was determined using an odor choice assay (Fig. 1A), which consists of a white plastic board as the arena, with a line drawn in the middle. Two lines were drawn on either side of the midline, at a distance that corresponds to the approximate length of the larvae, at the time of choice assay. Five drops of the control solution were added to a cotton ball that was placed at one end, 12cm away from the midline. Similarly, five drops of the odor solution were added to another cotton ball and placed at the other end. Each larva was aligned along the midline, and a white translucent plastic box, measuring 26.5cm X 9cm X 5cm was placed over the arena to cover and prevent any interfering visual and odor cues. A small piece of green tape was stuck at each (inner) end of the box so that the larvae, who are attracted to that color, move towards the ends of the arena and not towards the sides. Each larva (day 0, 5, 10 and 15) was given 4 minutes to walk to either side of the midline, except larvae at day 20, which were given 5 minutes. The choice made was noted at the end of the time interval by removing the plastic box that covered the arena. If the larva had crossed the line closer to the control side, it was noted to have chosen ‘control’. If the larva crossed the line closer to the odor side, it was noted to have chosen ‘odor’. If the larva had not crossed any of these lines, then it was noted to have made ‘no choice’.

**Fig 1.**
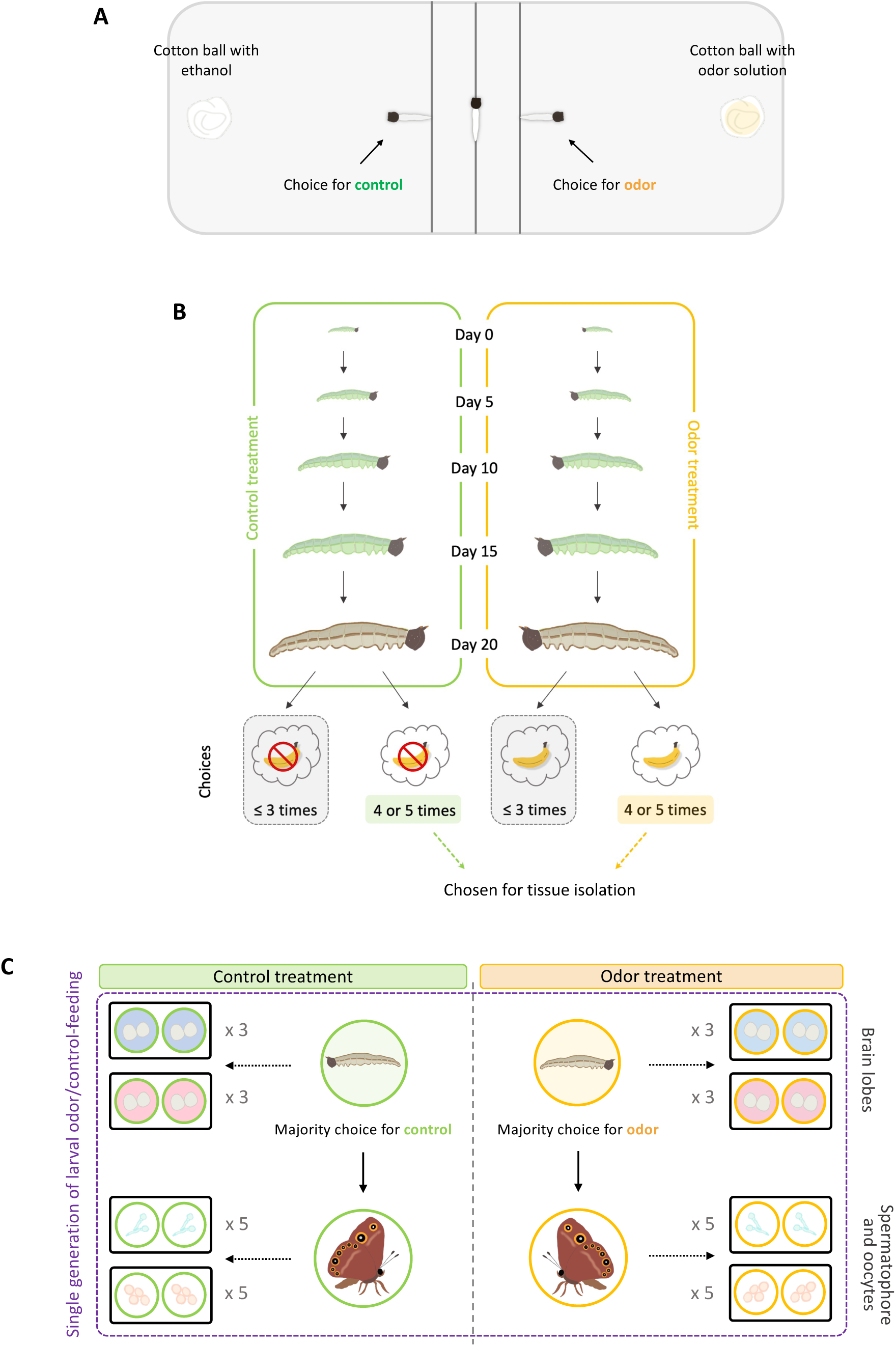
Experimental design. (**A**) *B. anynana* larvae were tested for either control or odor preference using the odor choice assay. (**B**) Larvae that showed a consistent preference for the treatment odor they were reared on (made at least 4 similar choices out of 5 attempts), were chosen for tissue isolation. (**C**) Male and female brains from selected larvae were isolated and collected separately. Spermatophores and oocytes were collected from selected adult butterflies. The numbers in grey denote the number of biological replicates.

### Selecting larvae that showed a majority preference towards odor or control

The larvae were tested again for their odor choices on days 5, 10, 15 and 20. Only larvae that consistently chose the odor they were reared on (either control or odor), for at least four out of the five times they were tested, were selected as the ones that showed a ‘majority preference’ (Fig. 1B). Twenty days after hatching, only those larvae (5^th^ instar) that chose either control or odor (depending on the treatment they reared on) at least four times out of the five times they were tested, were used for tissue isolation.

### Sample preparation

Brains were isolated from six 5^th^ instar larvae of each sex. The remaining larvae were allowed to pupate and eclose as adults. Butterflies were reared in cages and fed a diet of mashed banana. Oocytes (unfertilized eggs) were isolated from 10 female butterflies on the fifth day after they eclosed. Each male butterfly (ten in total) was allowed to mate with a naïve female butterfly that had not been exposed to any treatment as a larva. After mating, the spermatophore (a combination of male seminal fluids and sperm cells) that had been deposited in the female’s abdomen by the male butterfly during mating was dissected and isolated. Dissections were performed for both control and odor treatments (Fig. 1C). Tissue samples were collected in individually labelled Eppendorf tubes containing RNA later and stored at 4°C overnight. These tubes were then stored at −80°C until further processing.

### Nucleic acid extraction and sequencing

Each brain tissue group (for both control and odor treatments) had three biological replicates, with two brains of the same sex larvae pooled together in each replicate. Each oocyte and spermatophore group (for both control and odor treatments) had five biological replicates, with tissues from two butterflies pooled together in each replicate (Fig. 1C). Nucleic acid extractions were performed for each biological replicate of each tissue group using TRIzol.

Samples were quantified for total RNA, and sample integrity and purity using Nanodrop and Agilent 5400. NovogeneAIT Genomics Singapore Pte Ltd constructed mRNA libraries and sequenced 20 million 150 bp paired-end reads from each biological replicate using PE150 on Illumina platforms. Small RNA-sequencing was performed only on oocyte samples which had sufficient RNA left. The same company constructed small RNA libraries and sequenced 20 million 50 bp single-end reads from each biological replicate using SE50, on Illumina platforms.

### RNA-seq data curation and differential gene expression analysis

Raw RNA-seq data were trimmed and curated as described previously [21]. Adaptors were trimmed from the raw sequencing data using Trimmomatic 0.39 [22] to generate clean reads (options: PE ILLUMINACLIP: TruSeq3-PE.fa:2:30:10:8; true MAXINFO:40:0.2) (trimming statistics: Table S5.5). To choose longer reads over read correctness, this MAXINFO option was used as the raw reads quality was high. FastQC 0.11.9 was used for the quality control checks before and after adaptor trimming. To perform differential gene express analysis, the gene models were acquired from NCBI *B. anynana* v1.2 genome [23] (GCF_900239965.1); [24] with quasi-mapping mode (options: -validateMappings -seqBias -gcBias) was used to quantify all annotated transcripts in the clean reads (salmon mapping statistics: Table S1). R package tximport [25] to convert raw transcript counts that were imported in R studio to gene-level counts. DESeq2 [26] was used to normalize raw gene counts and to perform DE analysis.

### sRNA-seq data curation and miRNA analysis

Raw sRNA-seq data were trimmed and curated as described previously [21]. In brief, adaptors were trimmed from the raw reads using Trimmomatic 0.39 [27]. To annotate miRNAs, other types of small non-coding RNAs, such as rRNAs, tRNAs, snRNAs, and snoRNAs, were removed from the adaptor-trimmed clean reads using SortMeRNA 2.1b [28] (trimming statistics: Table S6). Next, clean reads between 17-25nt were used to annotate miRNAs.

A previous study annotated the first set of *B. anynana* miRNAs in the developing wings [21], but because there might exist novel miRNAs specific to new tissue types (oocytes), the *B. anynana* miRNAs were re-annotated with the new oocyte sequencing data using miRDeep2 0.1.3 [29]. sRNA reads were first trimmed and filtered between 17-25nt and then mapped to the NCBI *B. anynana* v1.2 genome using the mapper.pl module (options: -d -e -h -l 16 -m -q). Then, miRNA precursors were predicted using the miRDeep2.pl module using all the oocyte sequencing libraries with default settings. All previously annotated *B. anynana* mature miRNAs and precursors [21] were used as known miRNAs. All mature miRNAs of *H. Melpomene*, *B. mori*, and *D. melanogaster* from miRbase 22.1 [30] were used as closely related species.

The new miRNAs were first blasted against all miRNAs registered in miRbase 22.1 [30] and then against the previously annotated *B. anynana* miRNAs [21]. Those without a hit were considered novel. To generate a non-redundant set of mature miRNAs for naming purposes, all mature miRNAs were blasted against themselves to identify identical or highly similar sequences. The resulting conserved and novel miRNAs were named manually as described previously [21].

The expression of miRNAs was quantified using the quantifier.pl module of miRDeep2. Read counts were normalized, and differential expression analysis was performed using DESeq2 [26] in R Studio.

## Results

### Overall gene expression is clustered by tissue type

We examined overall gene expression differences across the sequenced libraries. The hierarchical clustering tree shows that the overall gene expression is clustered by tissue type (Fig. 2D). However, the control and odor treatments cannot be separated, showing they have similar expression profiles. The same can also be observed in the principal component analysis (PCA) of gene expression (Fig. 2E).

**Fig 2.**
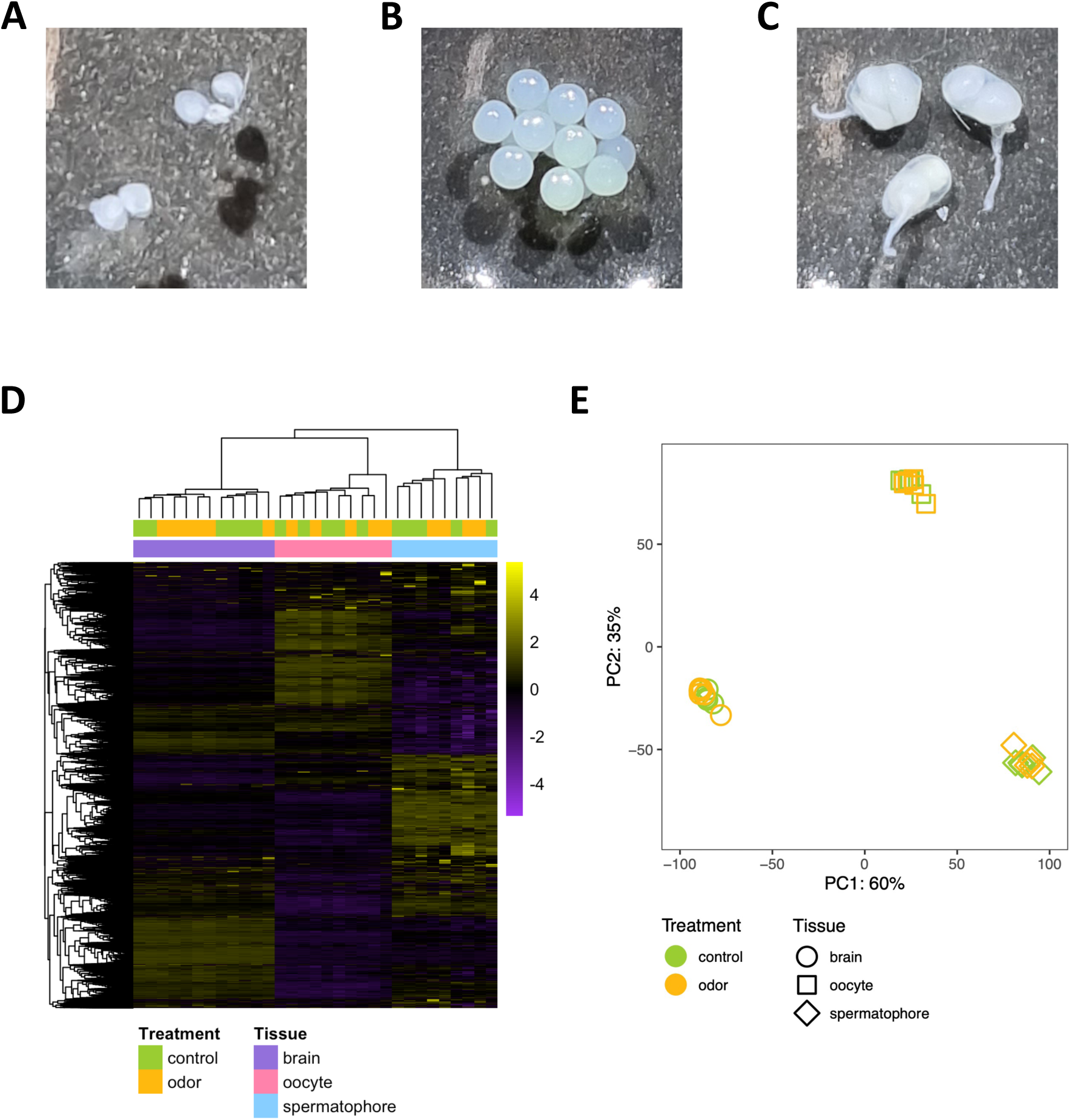
Overall gene expression is clustered by tissue type, but treatments are not separated. (**A**) Fifth instar larval brains (two are shown). (**B**) Oocytes from a single female adult butterfly. (**C**) Spermatophores (three are shown) collected immediately from a female’s body, after mating. (**D-E**) Genome-wide gene expression was evaluated using (**D**) hierarchical clustering heat map and (**E**) Principal Components Analysis (PCA).

### 83 genes were found to be differentially expressed between treatments

83 genes were found to be differentially expressed (*P*adj < 0.05) between treatment groups (Table 1). No DEGs were found between brains when the sexes were pooled. However, 14 genes were upregulated, and 7 were downregulated in the male brains of the odor treatment larvae relative to the control treatment. Similarly, 20 genes were upregulated and 28 were downregulated in female brains of the odor treatment larvae. In the oocyte comparison, 5 genes were upregulated and 5 were downregulated in the odor treatment. A single gene was upregulated, and 3 genes were downregulated in spermatophores of the odor treatment. A list of all the DEGs is provided in the supplementary information (male larval brains: Table S1; female larval brains: Table S2; adult butterfly oocytes: Table S3; adult butterfly spermatophores: Table S4).

**Table 1.** Several upregulated and downregulated genes in the tissues of larvae and adult butterflies of the odor group relative to the control group using DESeq2 analysis. These odor group larvae were fed (and preferred) the odor diet and the control group larvae were fed (and preferred) the control diet. *P*adj < 0.05.

| Tissues | # upregulated genes (in odor group) | # downregulated genes (in odor group) |
| --- | --- | --- |
| all brains | 0 | 0 |
| male brains | 14 | 7 |
| female brains | 20 | 28 |
| oocytes | 5 | 5 |
| spermatophores | 1 | 3 |

Out of the 83 genes that were found to be differentially expressed between treatments, 12 genes were either not annotated or identified as hypothetical or uncharacterized proteins. The function of each DEG was determined using literature searches. No single DEG was found to be shared among tissue types, such as between brains and germ cells, nor between male and female tissues (Fig. 3A).

**Fig 3.**
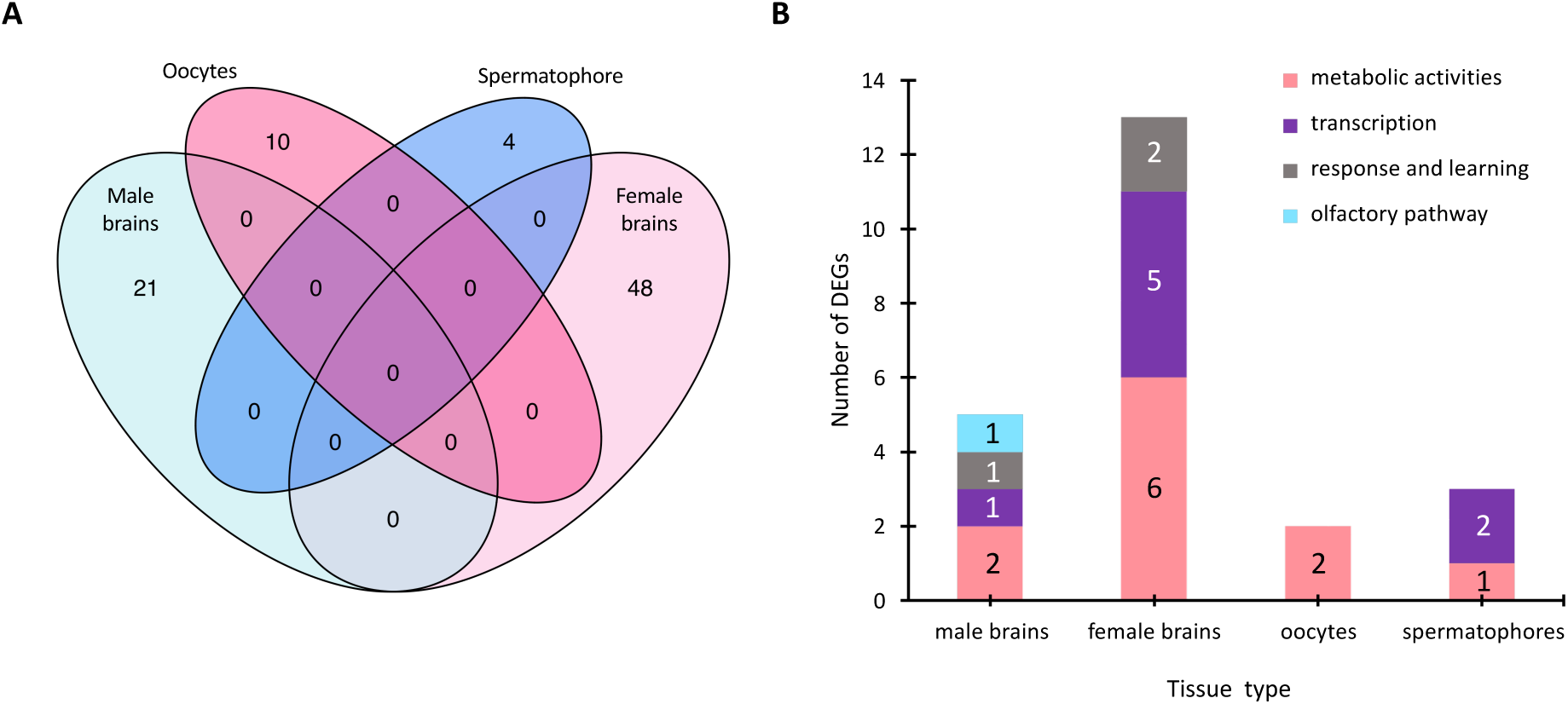
There are no shared DEGs among tissue types. **(A)** The male and female larval brains, oocytes and spermatophores do not share any common DEGs as depicted in this Venn Diagram. **(B)** Number of differentially expressed genes (mRNAs) for the comparison of treatments for each tissue sample. Each bar in the graph depicts the DEGs split by functional types.

### 23 differentially expressed mRNAs were found to have roles in various physiological functions of interest

Upon classifying the proteins, predicted from the DEGs, under general functions such as metabolic activities, transcription, response to different chemical stimuli and learning, and involvement in the olfactory pathway, 48 genes were disregarded owing to either low log2FoldChange (i.e., log2FC < 1 and >−1; except one gene due to its function being of interest - Gene ID: LOC112049058, log2FC = −0.7, *P*adj < 0.05), or functions that were not considered of interest to this study. The remaining 23 genes are highlighted in Fig. 3B and Table 2.

**Table 2.**
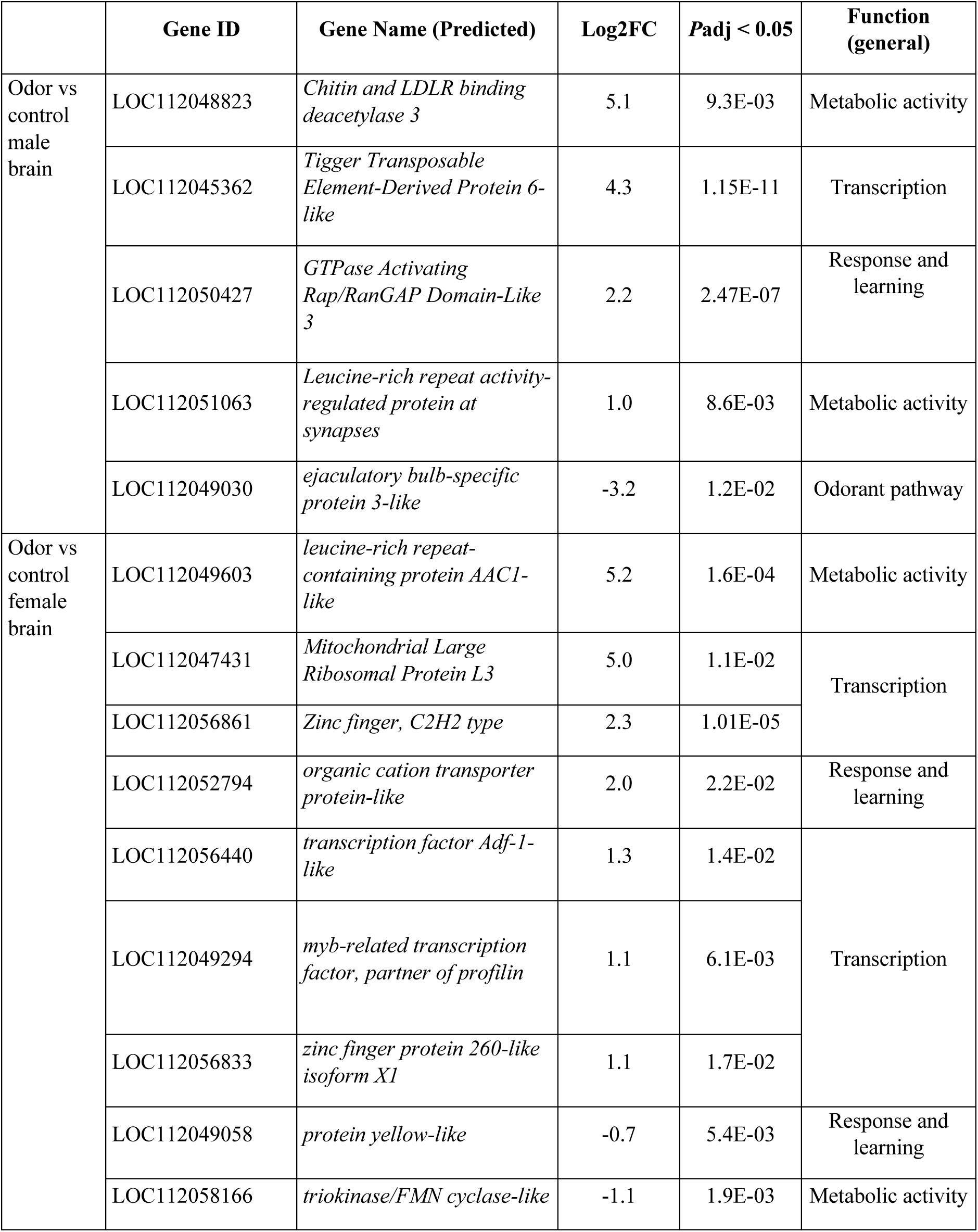
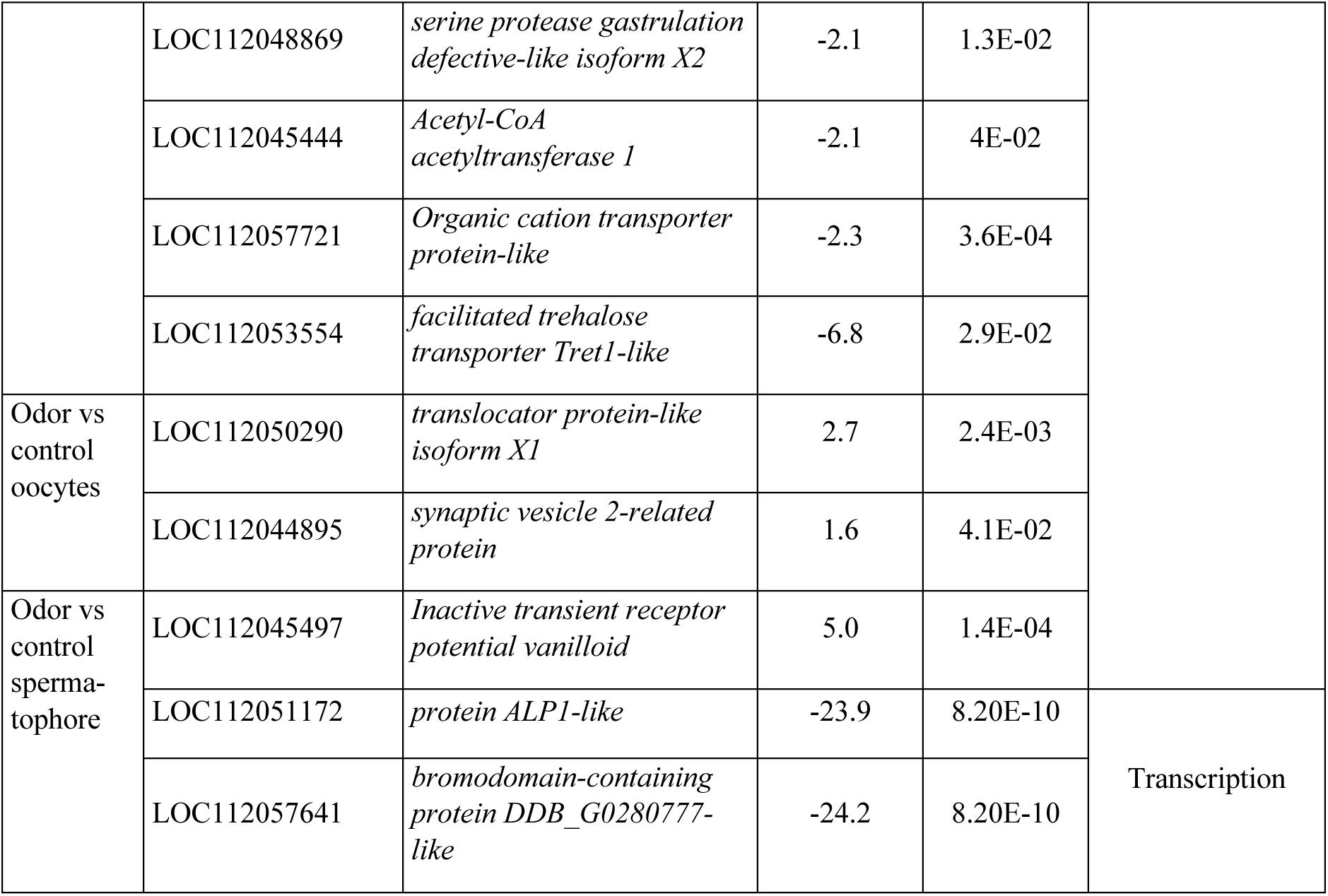
List of selected differentially expressed genes between tissues of odor and control groups. The ID of each DEG is highlighted with its predicted gene name, log2-fold change, adjusted *P*-value < 0.05, and its general function.

### No differentially expressed miRNAs were found in the oocytes of odor and control-fed female butterflies

The annotation of the sequenced miRNAs revealed 57 new miRNA precursors specific to oocytes (significant randfold *P*-value; miRDeep2 score > 3) and the full list of novel miRNAs are provided within the supplementary information (Table S7). Differential miRNA expression analysis across diet types was performed on these miRNAs, but control and odor treatment samples could not be separated using a PCA plot (Fig. 4), or using statistical methods. This suggests that diet does not impact the type and quantity of miRNAs observed across oocyte tissues.

**Fig 4.**
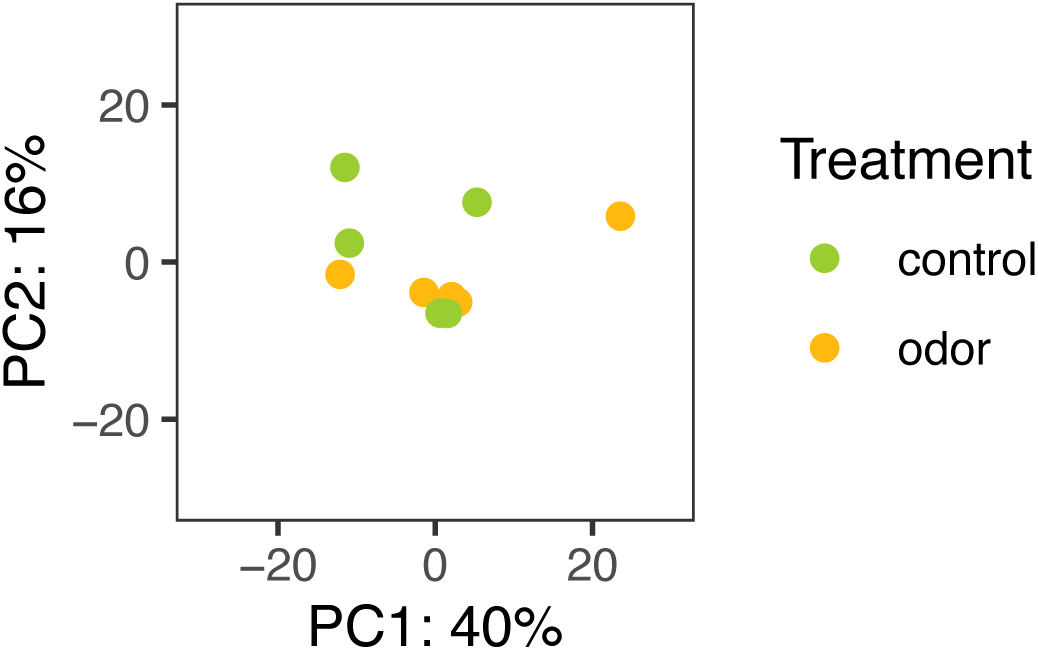
Control and odor treatments do not produce separated miRNA expression differences in female oocytes. The miRNA expression in oocytes of both control and odor treatments was evaluated using PCA.

## Discussion

In this study, we performed RNA sequencing on *B. anynana* larval brains, oocytes, and spermatophores of both control and odor treatments to investigate whether RNA epigenetic factors are induced during learning of a novel odor preference and transmitted to the next generation. We reared larvae on control and odor treatments and isolated tissue samples from larvae and butterflies that showed a majority larval preference for the treatment odor that they were reared on. After isolating and sequencing RNA from their brains and gonads, we found 23 mRNAs that might function in odor learning, three of which should be considered for further functional studies, but no good candidate that might function in odor preference inheritance.

Multiple mRNAs were found to be differentially expressed between the two odor treatment groups. Out of the 83 DEGs found, 40 of them were upregulated in the odor group and the remaining 43 were downregulated. Interestingly, not a single DEG was shared between tissue groups (Fig. 3A). This implies that the mRNAs that were differentially expressed during the odor exposure or involved in odor learning in the brain are not the same ones that are being passed down to the next generation. Perhaps, differentially expressed mRNAs found in brains of the odor-fed larvae might be involved in a pathway that induces different mRNAs in the germ tissues. However, this needs further testing. Moreover, the number of DEGs found in gonadal tissues between treatments (14 DEGs) is low in comparison to brain tissues (69 DEGs). As the DEGs in the gonads have no predicted roles in chemical sensing, but only in metabolism and transcription, they are probably of low significance to this study. Overall, there is little support for the roles of DEGs in being important epigenetic factors for the transfer of odor preferences from parents to offspring in this study.

Despite the lack of interesting candidate factors that might play a role in the inheritance of a learned odor preference, several genes were differentially expressed in male and female brains, and could be involved in the odor learning process, after odor feeding. In the male brain of the odor group, the *ejaculatory bulb-specific protein 3-like*, or *Pherokine-3* gene was downregulated relative to the control group (*Phk-3*, Gene ID: LOC112049030, log2FC = −3.2, *P*adj < 0.05). In *Drosophila*, *Phk-3* is an insect odorant/pheromone-binding protein (OBP), in addition to playing roles in metamorphosis and infection response [31,32]. In the biochemical pathway that detects various odorants, OBPs are the very first gene products that transport odorants to their membrane receptors in the antennal sensilla [33]. In a study on honeybees that were trained using lemon and strawberry odors coupled with a sugar reward, OBPs were both up or down-regulated (OBP3 – upregulated; OBP17 – downregulated), in their brains [34]. Thus, in the odor-fed larvae of the current experiment, *Phk-3* might have bound to the odorant molecules (IAA) and transported them to the sensory receptors of the olfactory nerve. It is important to note that in *Drosophila*, upon a fungal or bacterial infection, this *pherokine* was found to be more strongly expressed in the whole body of males than females [32]. In *B. anynana*, *Phk-3* was downregulated in the odor-fed male larval brains. Not much is known regarding the direct involvement of *Phk-3* in learning and memory. Thus performing functional studies to examine whether *Phk-3* is involved only in response to the odor molecules or also in memory formation of the odor experience might provide some insight into its specific role in the olfactory pathway.

A different set of DEGs were found to be differentially expressed in female larval brains between control and odor treatments. The *organic cation transporter protein-like* (CG7458, Gene ID: LOC112052794, log2FC = 2.0, *P*adj < 0.05) was found to be highly expressed in odor-fed female larval brains. It functions as a secondary active organic cation transporter in *Drosophila* [35]. This protein is orthologous to the mammalian transporters of the solute carrier family (SLC22) that are involved in remote sensing and signalling in the brain, in addition to mediating oxidative stress [36]. Hence, this gene could be involved in signal transduction in the brains of odor-fed larvae in this study. Another gene, *yellow*, was found to be downregulated in female larval brains of the odor treatment (Gene ID: LOC112049058, log2FC = −0.7, *P*adj < 0.05). *yellow* is transcribed into the major royal jelly protein (MRJP) and is of interest because in addition to its roles in eusocial behavior and caste specification, it is involved in neuronal signalling in honeybees [37,38], and in courtship behavior in *B. anynana* [39]. The genes *mrjp1* and *mrjp2* code for the major royal jelly proteins MRJP-1 and MRJP-2, which play roles in microbial infection defence and nutrition respectively, and were found to be upregulated in brains of honeybees trained to respond to sucrose odor [40]. In our experiment, however, *yellow* was downregulated in the odor-fed larval brains, and future functional studies might help us understand its role in the *B. anynana* olfactory or gustatory pathways.

Follow-up functional experiments could test the role of these DEGs in odor learning in *B. anynana*, however, some points deserve consideration. The DEGs of interest discussed above are sex-specific, thus knocking out these genes might either have a sex-specific effect, or might have some other function that has little to do with odor learning.

## Conclusion

The current work provides an entry point to the study of the molecular mechanisms of odor learning and memory formation in *B. anynana*, and the epigenetic factors that are involved in odor preference inheritance. Future studies should also sequence and analyse other small non-coding RNAs such as the piwi-interacting RNAs (piRNAs) as these are found abundantly in oocytes and ovaries of fruit flies, red flour beetles, and mammals [41–44]. In our study, however, due to technical difficulties in analyzing piRNA reads in the RStudio software, this analysis was not performed. Future studies could also examine alternative splicing of mRNA transcripts between the two treatment groups, for each tissue type, as they ultimately generate different proteins [21,45]. Also, single-cell RNA sequencing of the brain might capture cell-specific molecular responses, which may have been missed due to the bulk-sequencing approach used. Finally, (single-cell) DNA methylation should also be investigated, as some studies showed heritable methylation of specific loci by DNA methyltransferases (Dnmts) upon odor exposure [46,47].

## Supporting information

Electronic supplementary material - Additional file

## Acknowledgements

We thank the Fire Flies Health Farm and Greenology for providing corn plants.

## Funding

This study was supported by a Yale-NUS PhD scholarship to VG, the Ministry of Education (MOE) Singapore award MOE2018-T2-1-092, and the National Research Foundation, Singapore under its Investigatorship programme (award NRF-NRF105-2019-0006).

## Contributions

VG and AM conceptualized and designed this study. VG performed the experiments and collected the data. Both VG and ST analyzed the genomic data. ST analyzed the transcriptomic data. VG wrote the manuscript with input from all authors. All authors declare no conflict of interest.

## Ethics declaration

### Ethical approval

Not applicable.

### Consent to publish

Not applicable.

### Conflict of interest

The authors have no conflicts of interest to declare.

## Electronic supplementary material

### Additional file

**Table S1.** DEGs in the male larval brains of the odor group relative to the control group using DESeq2 analysis. **Table S2.** DEGs in the female larval brains of the odor group relative to the control group using DESeq2 analysis. **Table S3.** DEGs in the adult butterfly oocytes brains of the odor group relative to the control group using DESeq2 analysis. **Table S4.** DEGs in the adult butterfly spermatophores of the odor group relative to the control group using DESeq2 analysis. **Table S5.** RNA-seq data trimming and Salmon mapping statistics. **Table S6.** Oocyte sRNA-seq data trimming and Salmon mapping statistics. **Table S7.** Dataset containing the annotation of oocyte miRNAs in *B. anynana*.

## Notes

### Competing Interest Statement

The authors have declared no competing interest.

